# Lipid-protein interplay in dimerization of the juxtamembrane domains of epidermal growth factor receptor

**DOI:** 10.1101/197673

**Authors:** Ryo Maeda, Takeshi Sato, Kenji Okamoto, Masataka Yanagawa, Yasushi Sako

**Author notes:** Correspondance should be addressed to Y. S.

## Abstract

Transmembrane (TM) helix and juxtamembrane (JM) domains (TM-JM) bridge the extracellular and intracellular domains of single-pass membrane proteins, including epidermal growth factor receptor (EGFR). TM-JM dimerization plays a crucial role in regulation of EGFR kinase activity at the cytoplasmic side. Although the interaction of JM with membrane lipids is thought to be important to turn on EGF signaling and phosphorylation of Thr654 on JM leads to desensitization, the underlying kinetic mechanisms remain unclear. Especially, how Thr654 phosphorylation regulates EGFR activity is largely unknown. Here, combining single-pair FRET imaging and nanodisc techniques, we showed that phosphatidylinositol 4,5-bis phosphate (PIP_2_) facilitated JM dimerization effectively. We also found that Thr654 phosphorylation dissociated JM dimers in the membranes containing acidic lipids, suggesting that Thr654 phosphorylation electrostatically prevented the interaction with basic residues in JM and acidic lipids. Based on the single-molecule experiment, we clarified the kinetic pathways of monomer (inactive state) - dimer (active state) transition of JM domains and alteration in the pathways depending on the membrane lipid species and Thr654 phosphorylation.

---

Epidermal growth factor receptor (EGFR), the best studied receptor tyrosine kinase (RTK), plays an important role in regulating cell proliferation and differentiation^1,2^. EGFR consists of an extracellular domain that interacts with extracellular ligands, a single-pass transmembrane (TM) helix, a juxtamembrane (JM) domain, a cytoplasmic kinase domain that activates various signal cascades in the cell, and a C-terminal tail domain^3,4^. When ligands such as EGF bind to the extracellular domain of EGFR, conformational changes occur in the extracellular domain and the asymmetric dimer of intracellular kinase domains is formed^5^. This asymmetric dimerization enables one (activator) kinase domain to activate the other (receiver) kinase domain^6^, resulting in subsequent phosphorylation of tyrosine residues on the C-terminal tails of the activator and recruitment of intracellular signal proteins containing an SH2 and/or PTB domain, such as Shc, Grb2, etc.^7^.

Although the structures of most of the EGFR domains excluding the C-tail domain^5,6,8–10^ have been solved individually, the overall architecture remains unclear. In particular, the most important question regarding the regulation of EGFR kinase activity across the lipid bilayer, i.e., the correlation between conformational changes in the extracellular domain upon ligand binding and rearrangement of the intracellular kinase domain, is still controversial.

In the context of transmission of information across the membrane, the single-pass TM helix and the JM domain (TM-JM) play crucial roles in conformational coupling between the extracellular and cytoplasmic domains of EGFR^9,11,12^. Previous NMR studies and molecular dynamics (MD) simulations have suggested that the TM domain forms an α-helix dimer and its configuration is changed by the ligand-binding with the extracellular domains^9,13,14^. This information regarding conformational changes in the TM dimer is transmitted to the JM domains. The JM domain of EGFR consists of a JM-A (N-terminal half) region, which can form an antiparallel dimer and a JM-B (C-terminal half) region, which makes contact with the kinase domain. Both of the JM regions play important roles for stable formation of an asymmetrical kinase dimer conformation^11^. Especially, it is predicted that the antiparallel configuration of JM dimer leads to asymmetric dimerization and activation of the kinase domain^6,13^. The JM-A domain contains rich Lys or Arg residues, several of which are thought to interact with anionic lipid molecules of the plasma membrane^11^ and these lipid interactions are thought to be involved in antiparallel dimer formation^15^. However, the dynamic aspects of JM dimerization with regard to the interaction with membrane lipids remain unknown experimentally. For example, while previous studies have suggested that PIP_2_ interacts specifically with the basic residues on JM-A^15,16^, how PIP_2_ regulates binding of the JM domain to membranes and/or formation of JM dimers is not fully understood.

In addition to the activation process, a shutdown system is also crucial for signal regulation^17^. EGFR has many phosphorylation sites, mainly tyrosine residues in the kinase domain and C-terminal tail. In addition to these sites, Thr654 in the JM-A region is known to be phosphorylated by protein kinase C (PKC)^18^. Previous biochemical studies have suggested that kinase activity of EGFR is decreased when this Thr654 is phosphorylated^19^. Furthermore, Thr654 is known to be a calmodulin-binding site and phosphorylation of Thr654 suppresses the binding^18^, resulting in inhibition of EGFR autophosphorylation^20^ and internalization^21^. Until recently, despite the important role of Thr654 phosphorylation, little is known about the structural insights into the phosphorylated states.

Here, we applied single-pair fluorescence resonance energy transfer (spFRET) analysis using total internal reflection fluorescence (TIRF) microscopy to investigate the conformational regulation mechanisms by membrane lipid molecules and Thr654 phosphorylation. We have demonstrated previously that single-molecule analysis is a powerful tool to directly monitor the physiologically relevant conformational changes in molecules, such as rhodopsin, which is a G protein-coupled receptor^22^ and DNA Holliday junction^23,24^. To examine the effects of lipid components on dimer conformation of TM-JM domain of EGFR, TM-JM peptides were synthesized, conjugated with fluorophores, and introduced into nanodiscs. Nanodiscs are small (10-12 nm in diameter) membrane lipid bilayers consisting of membrane scaffold proteins (MSPs) and lipid molecules^25,26^. By changing the ratios of receptor and MSP and/or lipid species, the number of receptor molecules and membrane lipid composition can be regulated in the nanodiscs^27,28^. Here, by a combination of spFRET analysis and nanodisc systems, we assessed the conformational dynamics of formation/deformation of JM dimer structure and revealed the conformational regulation mechanism depending on the membrane lipid composition, paying attention to the effects of acidic lipids. Furthermore, we compared the dynamics of dimer formation of the JM regions in non-phosphorylated and Thr654 phosphorylated states, and discuss how the phosphorylation of this site leads to inactive conformations of EGFR.

## Results

### Incorporation of TM-JM peptides into nanodiscs

Synthesized peptides of TM-JM region of EGFR were prepared as described previously and labeled with the fluorophore Cy3 (TMJM-Cy3) or Cy5 (TMJM-Cy5)^15,29^. The peptides were mixed with purified membrane scaffold proteins (MSPs) and lipid molecules, here POPC (PC) and POPS (PS), in the ratio of TM-JM:MSP:lipids = 1:1:120 in molar ratio, and reconstituted into nanodisc structures by removing detergents (Fig. 1a). The nanodisc particles were isolated by gel filtration size exclusion chromatography (Fig. 1c), and the peak fractions of nanodiscs were collected and further characterized. Negative stain electron microscopy clearly showed that uniform nanodisc particles were formed and purified successfully (Fig. 1b). Fluorescence emission spectroscopy of the nanodisc fraction prepared using a 1:1 mixture of Cy3- and Cy5-labeled peptides showed that both peptides were reconstituted into the same nanodisc fractions (Fig. 1e). In addition, as shown in Fig. 1e, Cy5 emission caused by FRET from Cy3 was observed, indicating that TMJM-Cy3 and TMJM-Cy5 peptides in the same nanodiscs form heterodimers. Synthesized pTM-JM peptides, in which Thr654 was phosphorylated, were also reconstituted into nanodiscs (Fig. 1d and 1f). As shown in Fig. 1c and d, the composition of lipid molecules consisting of nanodisc structures (PC:PS = 10:0, 10:3, and 3:10) has little effect on the incorporation of TM-JM peptides. Sensitized emission of Cy5 caused by FRET was observed in all of the membrane compositions. The peak fractions of TM-JM nanodiscs (Fig. 1c, d) were applied to the following single-molecule measurements.

**Figure 1.**
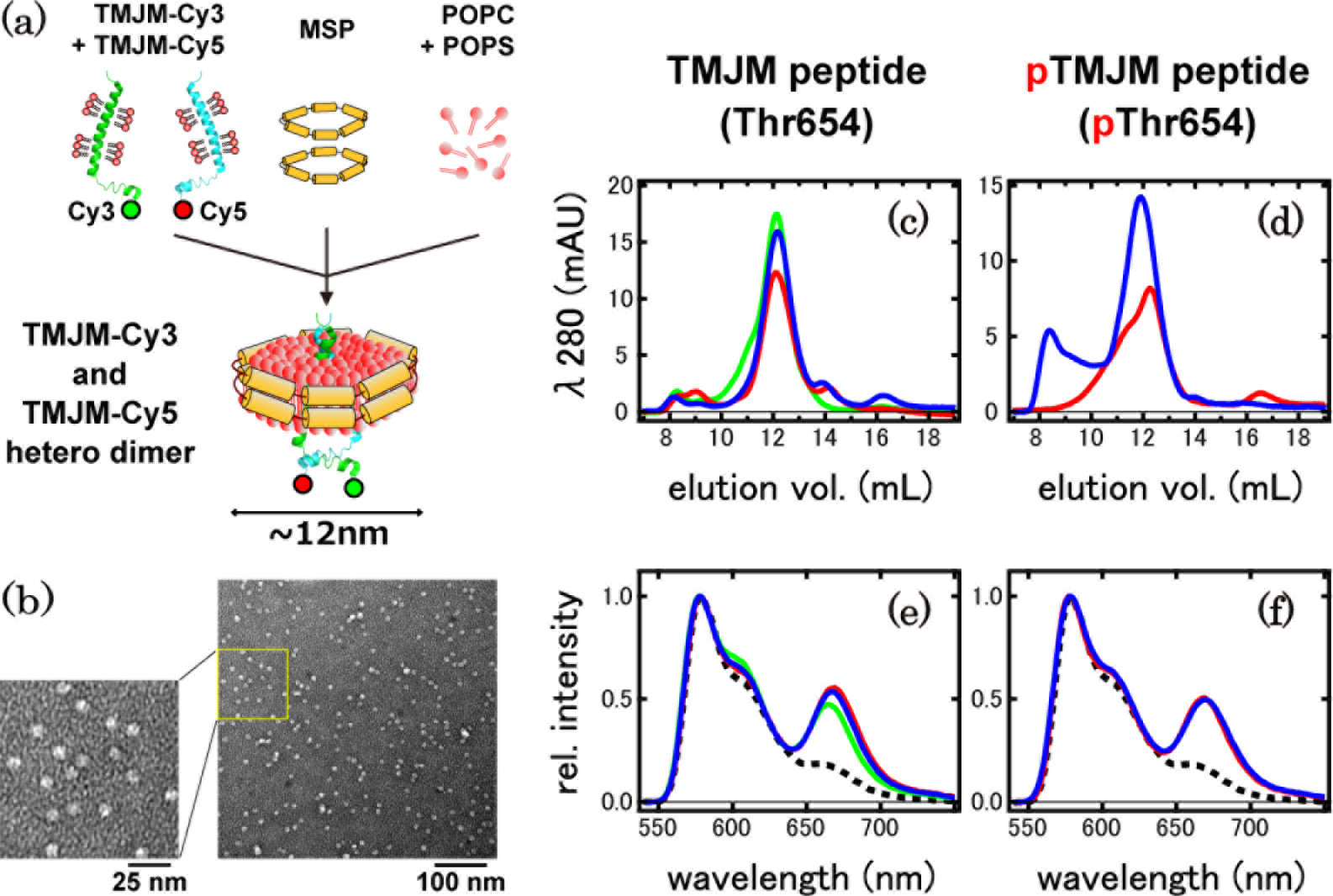
Reconstitution of TM-JM peptides into nanodiscs. (a) Schematic diagram of the preparation of nanodisc samples. (b) Electron microscopic images of nanodisc particles. The sample was acquired from the fraction of elution volume = 12 mL in (c) and stained with 2% samarium acetate. (left) An enlarged image of the boxed yellow region. (c,d) Gel filtration chromatography of nanodiscs containing (c) non-phosphorylated TMJM-Cy3 and TMJM-Cy5 peptides or (d) phosphorylated TMJM-Cy3 and TMJM-Cy5 peptides. Line colors indicate lipid compositions; PC:PC = 10:0 (red), PC:PS = 10:3 (blue), and PC:PS = 3:10 (green). (e,f) Bulk fluorescence spectroscopy of nanodisc samples containing (e) non-phosphorylated or (f) phosphorylated TMJM-Cy3 and TMJM-Cy5 peptides. Dashed lines show the summation of fluorescence of Cy3 and Cy5 dyes with excitation at 540 nm.

### Single-pair FRET (spFRET) imaging

The TM-JM nanodiscs were immobilized onto the glass surfaces covered by polyethylene glycol (PEG) as described previously^30,31^ using the association between NeutrAvidin on biotinylated PEG (b-PEG) and biotinylated anti-His6-tag antibody bound to His-tagged MSPs (Fig. S1a). To confirm the blocking effects on nonspecific binding, we compared single-molecule images, and one contained b-PEG and the other did not. Figures S1b and c clearly showed that the number of fluorescent spots was significantly increased on b-PEG-containing glass compared to that lacking b-PEG, indicating that TM-JM nanodiscs were successfully anchored on the glass surfaces via MSP molecules at the edge of the nanodiscs.

TMJM-Cy3 and TMJM-Cy5 heterodimeric peptides in nanodiscs were illuminated with a 532 nm laser (Fig. 2a and Movie S1). Some fluorescent spots showed Cy5 fluorescence derived from the occurrence of FRET by proximity between Cy3 and Cy5 labeled on the C-terminus of JM-A domains. Intensity histograms constructed from fluorescent spots showing Cy5 emission demonstrated that most of the FRET spots contained single TMJM-Cy3 and TMJM-Cy5 molecules (Fig. S2a - d).

**Figure 2.**
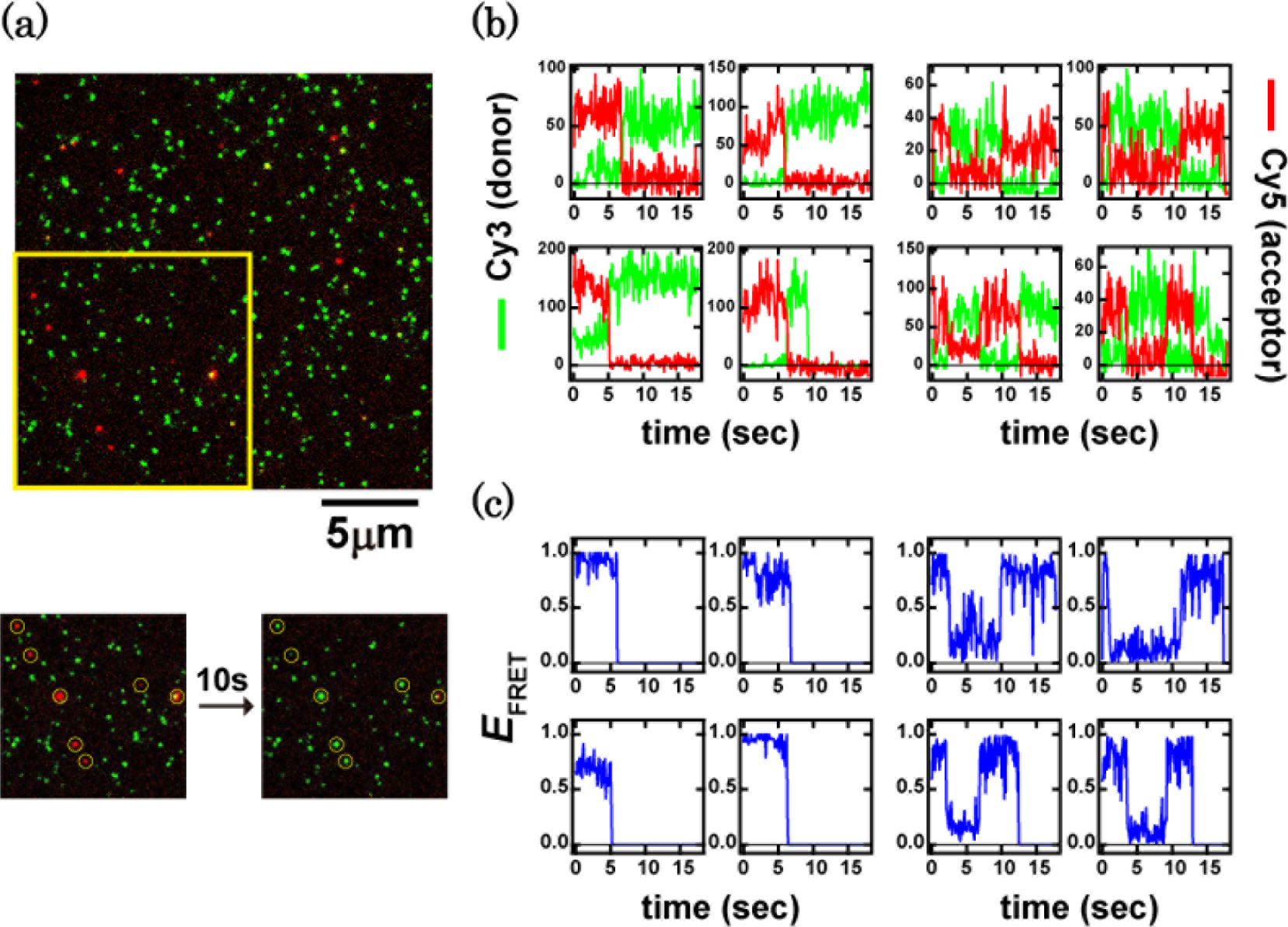
Single-pair FRET observation of TMJM-Cy3 and TMJM-Cy5 dimers. (a) (upper) Typical TIRF image of nanodisc samples containing TMJM-Cy3 and TMJM-Cy5. The lipid composition was PC:PS = 10:3. Cy3 was excited with a 532 nm laser, and fluorescence of Cy3 (green) and Cy5 (red) was recorded at 100 ms/frame. (lower) Typical images of fluorescent spots before and after Cy5 photobleaching. (b) Representative fluorescence trajectories of Cy3 (green) and Cy5 (red). (c) FRET efficiency trajectories of (b). FRET efficiency *E*_FRET_ was calculated as described in Material and Methods.

Typical fluorescence trajectories derived from TMJM-Cy3 and TMJM-Cy5 in the lipid composition of PC:PS=10:3, which was relatively similar to mammalian cell membranes^32^, are shown in Fig. 2b. FRET efficiency *E*_FRET_ was calculated from these traces (Fig. 2c, detailed in Material and Methods). In some nanodiscs, TMJM-Cy3 and TMJM-Cy5 peptides showed stable high FRET signals before Cy5 photobleaching (Fig. 2b and 2c, left). As the Förster radius *R*_0_ between Cy3 and Cy5 is 5.6 nm^33^, this result suggests that TM-JM peptides form a stable dimer conformation, which positions the C-terminus of JM-A domains in proximity (hereafter, we refer to this conformation as JM dimer). However, other populations of spots showed transition of FRET efficiency before Cy5 photobleaching (Fig. 2b and 2c, right). The transitions to low FRET efficiency states also suggested that dissociation of JM dimer occurred occasionally.

When TM-JM peptides were labeled at the N-terminus with Cy3 or Cy5, most of the FRET spots showed high Cy5 fluorescence as well as extremely low Cy3 fluorescence intensity (Fig. S3a), providing a FRET distribution with a large population of relatively high FRET efficiency (*E*_FRET_ > 0.5) (Fig. S3b, blue). Moreover, most (~95%) of the peptide dimers contained a high FRET efficiency state in the trajectories (Fig. S3b, red), indicating that TM-JM peptides are in contact with each other at TM domains in nanodisc membranes in the parallel orientation and N-termini of the peptides are in close proximity (Fig. S3b, inset).

### Lipid and phosphorylation effects on JM dimerization

To examine the effects of lipid molecules on JM dimer formation, FRET efficiency distributions were determined with various nanodisc lipid compositions (Fig. 3a-d). With a lipid composition of PC:PS = 10:0, both the low and high FRET efficiency states were detected with a broad distribution around the intermediate FRET efficiency (Fig. 3a, red line). It is plausible that the low and high FRET populations represent the conformations in which the JM-A domains of each protomer are dissociated (described as JM-monomer) and associated (JM-dimer), respectively. On the other hand, there was a wide range of distributions of FRET states between the low and high FRET states, suggesting that there are many conformational substrates between JM-monomer and JM-dimer. For convenience, we divided the distributions into three populations—JM-monomer (0 ≤ *E*_*FRET*_ < 0.33), JM-dimer (0.67 ≤ *E*_*FRET*_), and intermediate populations (0.33 ≤ *E*_*FRET*_ < 0.67; JM-intermediate)—and compared the fractions in each population (Fig. 3g-i). Addition of PS (PC:PS = 10:3, Fig. 3a, blue line) decreased the fraction of JM-monomer but increased that of JM-dimer (Fig. 3g), indicating that PS stabilized the JM-A dimer conformation. However, in membranes containing larger amounts of PS (PC:PS = 3:10), the population of JM-dimer was decreased to a similar level as shown in PC:PS = 10:0 (Fig. 3b). In bulk fluorescence spectroscopy, Cy5 emission caused by FRET differed slightly among all of the membrane compositions, which was inconsistent with the spFRET measurements (Fig. 1e,f). This discrepancy was likely due to the low probability of incorporation of TM-JM peptides with the TMJM-Cy3 and TMJM-Cy5 heterodimeric combination into nanodiscs, as shown in Fig. 2a.

**Figure 3.**
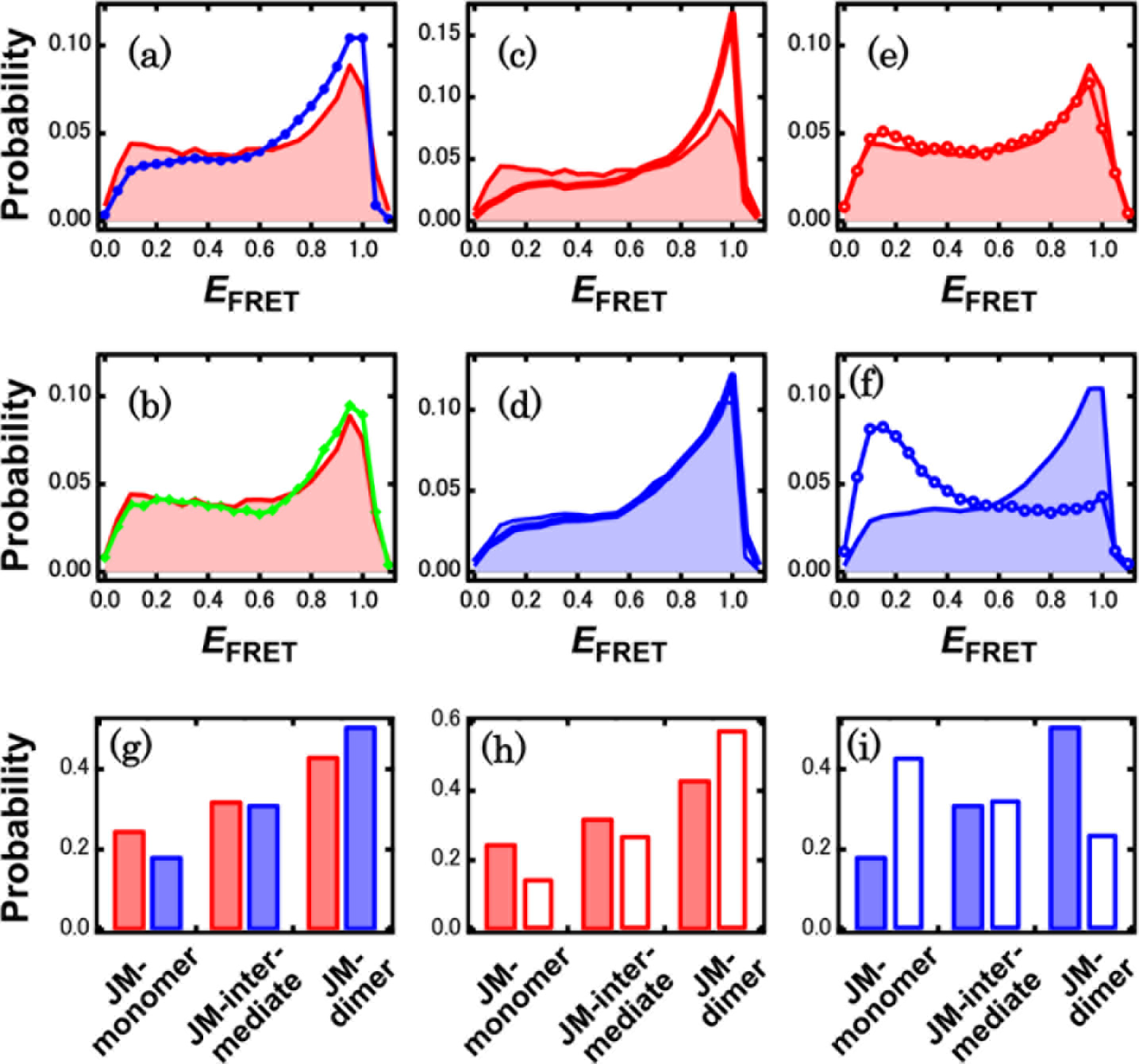
FRET efficiency distributions of TMJM-Cy3 and TMJM-Cy5 dimer. (a,b) FRET efficiency (*E*_FRET_) distributions of nanodiscs containing non-phosphorylated TMJM-Cy3 and TMJM-Cy5 peptides with lipid compositions of PC:PS = 10:0 (red), PC:PS = 10:3 (blue), and PC:PS = 3:10 (green). FRET efficiency distributions of nanodiscs containing non-phosphorylated TMJM-Cy3 and TMJM-Cy5 peptides in (c) PC:PS = 10:0 or (d) PC:PS = 10:3 membranes before (filled regions) or after (lines) addition of PIP_2_. FRET efficiency distributions of nanodiscs containing Thr654 phosphorylated TMJM-Cy3 and TMJM-Cy5 peptides in (e) PC:PS = 10:0 (red circle) or (f) PC:PS = 10:3 (blue circle) membranes. (g-i) Comparison of the fractions of JM-monomer (0 ≤ *E*_*FRET*_ < 0.33), JM-intermediate (0.33 ≤ *E*_*FRET*_< 0.67), and JM-dimer (0.67 ≤ *E*_*FRET*_ ≤ 1) states. The values were calculated by summation of the corresponding distributions in (a-c). The number of total frames utilized for construction of the distributions was 56,000-175,000 in each condition.

Addition of negatively charged lipid PIP_2_ to the nanodiscs with PC:PS = 10:0 markedly shifted the population of FRET efficiency toward high FRET states (Fig. 3c). This increase was larger than in the case of PS (Fig. 3g,h), suggesting that PIP_2_ strongly stabilized the dimer conformation of JM-A domains. On the other hand, addition of PIP_2_ to the nanodiscs containing PC:PS = 10:3 showed little effect on the FRET distribution (Fig. 3d), suggesting that the roles of PS and PIP_2_ in JM-A dimer formation partially conflicted.

Next, we examined the effects of Thr654 phosphorylation on TM-JM dimerization. As shown in Fig. 3f and i (PC:PS = 10:3), phosphorylation of Thr654 markedly altered the distribution of FRET efficiency, in which JM-dimer decreased and JM-monomer increased. These results were consistent with the report that phosphorylation of Thr654 leads to inactivation of EGFR^19^. It should be noted that phosphorylation of Thr654 did not markedly change the FRET distribution under a lipid composition of PC:PS = 10:0 (Fig. 3e), suggesting that phosphorylation of Thr654 electrostatically interferes with the interaction between negatively charged lipid PS and basic residues in JM, and dissociation from the membrane is coupled with the dissociation of JM dimers (see Discussion).

### Bayesian change point detection in the spFRET trajectories

We next analyzed the transition of FRET efficiency states to dynamically examine the conformational changes of TM-JM dimers. As shown in Fig. 2b and c, transitions between different FRET efficiency states occurred in a stepwise manner. Therefore, we examined the conformational state transitions using the modified Bayesian change point detection (BCPD) method^34^. The parameters were optimized for the detection method (Fig. S4), and we obtained change points in each condition (Fig. S5). All of the identified *E*_FRET_ values were collected and the FRET distributions were reconstructed by taking the time-weighted average within a bin size 0.05 (Fig. S5b-d). As shown in Fig. S5b and c, addition of PS or PIP_2_ increased the probability of high FRET states. On the other hand, when Thr654 was phosphorylated, the distribution was dominated by the population of low FRET states (Fig. S5d). These results were consistent with previous distributions constructed by accumulating all of the time points (Fig. 3a,c, and f), confirming that our Bayesian methods for detection of the FRET change points are reasonable.

The FRET distributions were divided into three groups according to *E*_FRET_ value, i.e., JM-monomer, JM-intermediate, and JM-dimer states, as shown in Fig. 3, and we examined the lifetime distributions of each state. All of the distributions were well fitted with single exponential functions, and their lifetimes were considerably shorter than the observation time limited by photobleaching (Fig. S6). Thus, the degree of kinetic heterogeneity among the substrates contained in each three group was not large.

### Transition of TM-JM dimer conformation

To explore the transition pathway between different FRET states, we constructed two dimensional transition maps plotting the frequency of transitions as a function of the initial (before transition) and final (after transition) *E*_FRET_ values detected using the BCPD algorithm, as described above (flow map; Fig. S7a-d). All flow maps were almost symmetric against the diagonal line of the *E*_FRET_ values (dotted line), indicating that detailed balance between the forward and backward transitions was observed among all conformational states under all experimental conditions.

Next, we constructed two-dimensional transition maps plotting the transition rate constants (rate constant map; Fig. 4a-d). These maps showed a rate constant, calculated by dividing the number of transition events to the final FRET states by the integrated lifetime of the initial FRET states. The maps were asymmetric to the diagonal lines due to differences in the forward and backward reaction rate constants (Fig. 4a, c). Under conditions with high dimer fractions, transitions from lower to higher FRET states were faster than the reverse transitions, as expected. To visualize the effects of changes in the membrane environment or phosphorylation of Thr654, subtractions of the rate constant maps were constructed (Fig. 4e-g). As shown in Fig. 4e, addition of PS increased the transition rate constant from low to high FRET populations (red circles), while decreasing those from low to intermediate and from intermediate to high FRET populations (blue circles). Although addition of PIP_2_ increased the transition rate constant from low to high FRET populations, similar to PS (red circle), PIP_2_ also increased the rate constants from low to intermediate and from intermediate to high FRET populations (green circles), which was the opposite of the effect of PS (Fig. 4f). These roles of PIP_2_ effectively shifted the conformational equilibrium toward dimerization of JM-A domains (Fig. 3g,h; see Discussion). When Thr654 was phosphorylated, transition rate constants from higher to lower FRET populations were increased, whereas those from lower to higher FRET populations were decreased (Fig. 4g). Thus, phosphorylation of Thr654 shifted the equilibrium largely toward dissociation of JM-A domains (Fig. 3i; see Discussion).

**Figure 4.**
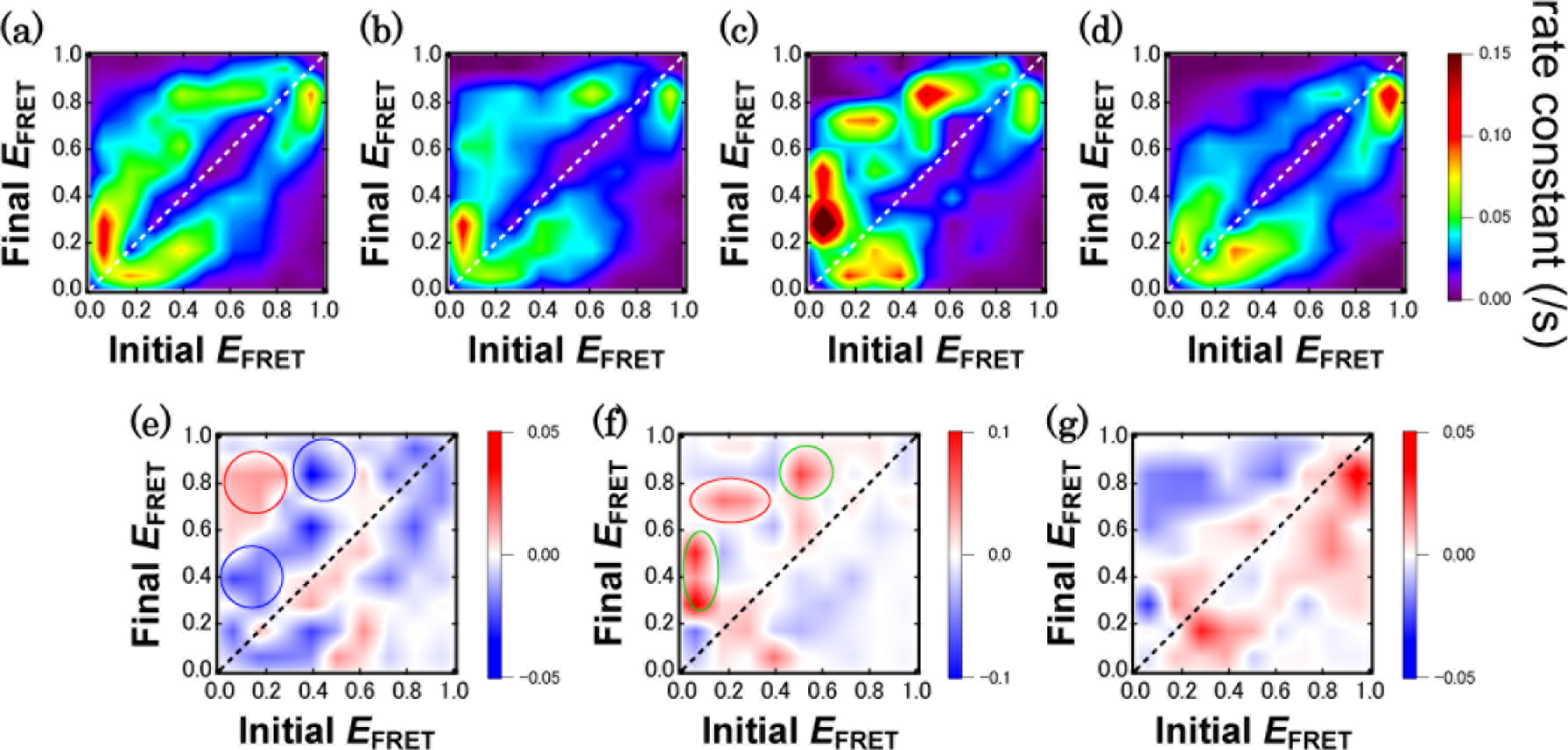
Two-dimensional rate constant maps. (a-d) Two-dimensional transition maps plotting the transition rate constants between the initial and final FRET efficiency values detected using the Bayesian change point detection algorithm. (a) Non-phosphorylated peptides in PC:PS = 10:0 membranes. (b) Non-phosphorylated peptides in PC:PS = 10:3 membranes. (c) Non-phosphorylated peptides in PC:PS = 10:0 membranes after addition of PIP_2_. (d) Thr654 phosphorylated peptides in PC:PS = 10:3 membranes. FRET efficiencies were divided into 9 × 9 cells in calculation and the surface plots were generated using Igor software. (e-g) Subtraction of 2D state transition flow maps which focused on (e) the effect of PS; (b) - (a), (f) the effect of PIP_2_; (c) - (a), and (g) the effect of Thr654 phosphorylation; (d) - (b).

## Discussion

Here, we studied the effects of membrane lipid composition and JM phosphorylation on the TM-JM dimer conformations of EGFR by applying nanodisc techniques to spFRET measurements. Synthesized fluorescent peptides of TM and JM-A regions of EGFR were reconstituted into nanodisc structures in which the compositions of lipid molecules were altered. The peptides formed dimers at the TM domain with a high probability, and interaction dynamics between the JM-A region were detected from the FRET trajectories between the fluorophores at the C-terminus of the peptides. The JM-A domain is rich in Lys and Arg residues, and it is widely accepted that several of these basic residues interact with acidic lipids in the plasma membrane^11^. On the other hand, an antiparallel helix dimer structure of JM-A was found in previous NMR and MD simulation studies^9,13^. In the antiparallel dimer structures, the LR_656_R_657_LL motif in JM-A stabilizes the dimer conformation by hydrophobic interactions between Leu residues. Association with the acidic membrane lipids would be coupled with the antiparallel dimer formation of the JM-A region to regulate EGFR activity.

In our study, the TM-JM peptides showed a large fraction of high FRET states at the C-terminus of JM in the presence of negatively charged lipid PS (PC:PS = 10:3), indicating that JM protomers efficiently formed dimer conformations, which are likely to be in the antiparallel helix conformation. It has been reported that the TM-JM domain of EGFR reconstructed in bicelles with a lipid composition of PC:PS = 10:3 forms an antiparallel dimer at the JM-A regions with a distance of about 20 Å between the glutamate residues on the C-termini of the protomers^9^. Such close proximity will result in extremely high FRET efficiency, as detected in our experiments (Fig. 2c). Interestingly, PIP_2_, a key element for EGFR activation^35^, increased FRET efficiency more effectively than PS (Fig. 3a and c), indicating that PIP_2_ interacts with JM-A regions and stabilizes the antiparallel dimer structure more specifically. In contrast to the effects of negatively charged lipids, phosphorylation of Thr654, which is involved in deactivation of EGFR, decreased the probability of high FRET states and increased the low FRET states. This result indicated that phosphorylation of Thr654 leads to dissociation of the JM-A dimer structures.

One of the advantages of spFRET measurements is to allow monitoring of transitions between specific FRET states for kinetic analysis. We obtained FRET time course trajectories and detected change points of FRET efficiency to construct 2D maps of the transition rate constants between the FRET states. Comparison of the transition rate constant maps between membranes containing and lacking PS indicated that PS increases the rate constant from low to high FRET populations (Fig. 4e and 5a). As the presence of PS increased the probability of JM-dimer but decreased that of JM-monomer (Fig. 3g), PS was thought to have two effects, i.e., lowering the free energy of JM-dimer and lowering the activation energy from JM-monomer to JM-dimer (Fig. 4a). Assuming that the negatively charged PS increases the affinity between the membrane and the positively charged JM-A region, PS induced membrane association and dimerization of JM-A cooperatively. On the other hand, PS slightly stabilized the JM-intermediate states, as indicated by the decreases in transition rate constants from low to intermediate and intermediate to high FRET populations (Fig. 4e). It is possible that associations between PS and some of the positively charged amino acid residues trap the JM region in configurations that prevent dimerization. This may be why PS-induced dimerization was lost at higher PS concentrations (Fig. 3b).

Addition of PIP_2_ to the nanodiscs markedly increased the probability of the JM-dimer state but decreased that of the JM-monomer state (Fig. 3c,h), i.e., PIP_2_ significantly lowered the free energy of JM-dimer relative to PS as well as increasing that of JM-monomer (Fig. 4b). Moreover, in contrast to PS, addition of PIP_2_ increased the transition rate constant from low to intermediate FRET populations as well as low to high FRET populations (Fig. 4f and 5b). PIP_2_ also increased the transition rate constant from intermediate to high FRET populations. Thus, in contrast to PS, PIP_2_ induced JM dimerization without stabilizing JM-intermediate states, indicating that interaction with the JM region is not identical between PS and PIP_2_. Previous fluorescence spectroscopic studies^15,29^ and MD simulations^16^ indicated that PIP_2_ is distributed around the dimer more proximately than PS, interacting with multiple basic residues in the JM-A regions. Furthermore, it was also suggested that distance from the membrane to the LRRLL motif is greater for PIP_2_ in the PC/PS/PIP_2_ membrane than for PS in the PC/PS membrane^16^, which was thought to be essential for kinase activation. These differences in the interaction with the JM-A region between PS and PIP_2_ must be related to the differences in JM dimerization kinetics identified in this study. We anticipate that PIP_2_ but not PS would effectively push aside the LRRLL motif and maintain an appropriate distance from the membrane surface, inducing the appropriate orientation for antiparallel helical dimer formation (Fig. 5b).

**Figure 5.**
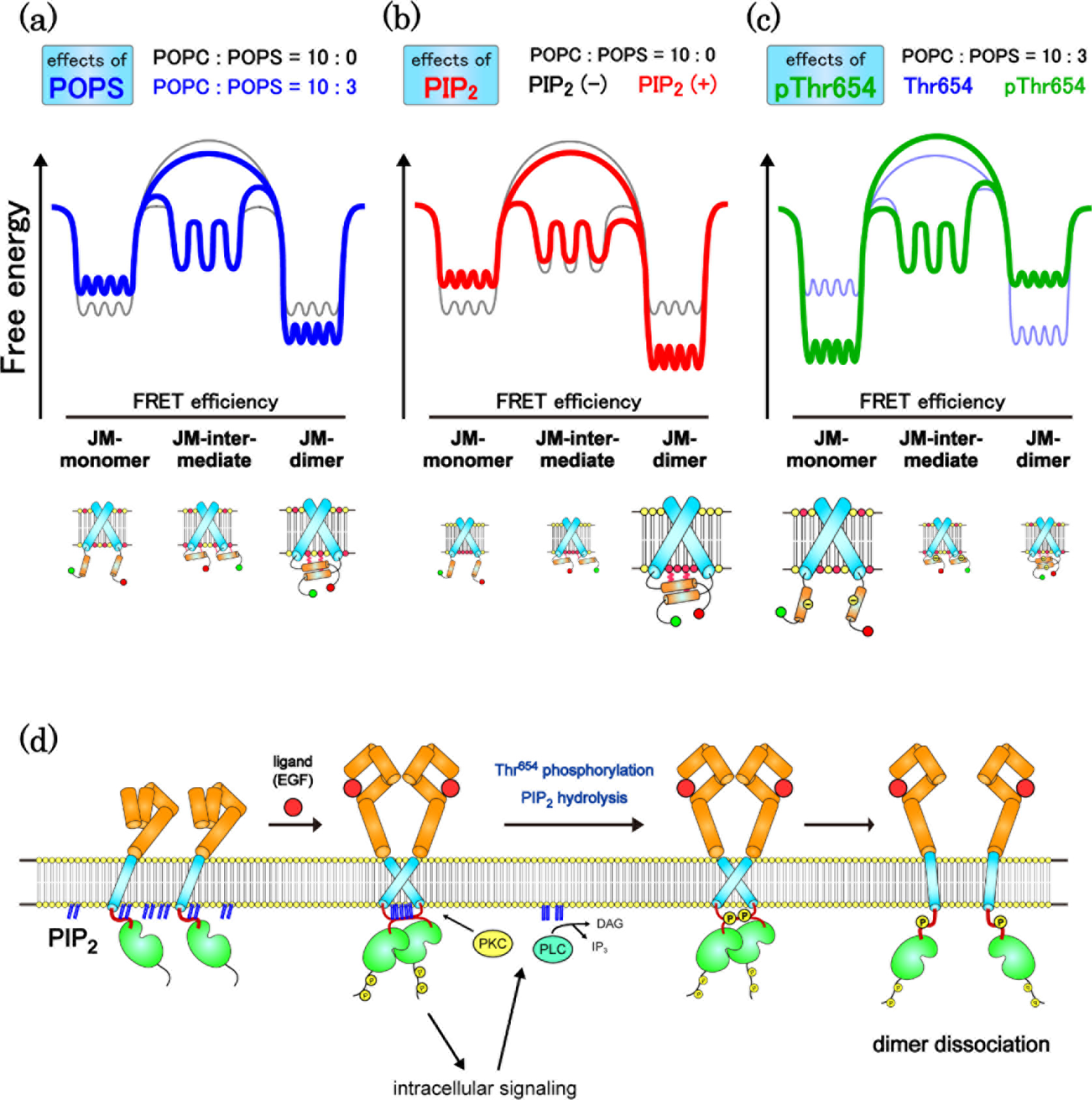
Effects of JM-lipid interactions on the TM-JM dimer formation. Onedimensional projections of the energy landscape of TM-JM dimer conformation in three comparisons: (a) Effect of the presence of PS in the membrane (blue). (b) Effect of addition of PIP_2_ to the membrane (red). (c) Effect of phosphorylation of Thr654 (green). See text for details. (d) A schematic image of the regulation of EGFR activity by JM-PIP_2_ interactions.

Limited structural information is available regarding the Thr654-phosphorylated TM-JM. In the present study, Thr654 phosphorylation increased JM-monomer in PC:PS = 10:3 membrane but not in PC:PS = 10:0 membrane (Fig. 3e,f). In the PC:PS = 10:3 membrane, in contrast to PIP_2_, Thr654 phosphorylation markedly decreased and increased the free energy of JM-monomer and JM-dimer, respectively, markedly shifting the conformational equilibrium toward dissociation of the JM dimer (Fig. 4c). In addition, Thr654 phosphorylation decreased the transition rate constants from lower to higher FRET populations, but increased the rate constants from higher to lower FRET populations (Fig. 4g and 5c). The dependence on PS concentration suggests that the phosphorylation electrostatically interferes with the association between JM and acidic membrane lipids, but does not directly inhibit dimerization within JM protomers. Dissociation from the membrane would result in dissociation of the dimeric structures. This mechanism is consistent with the effects of acidic lipids, which induce JM dimerization by increasing the affinity of JM with the membrane. Thr654 is adjacent to Arg656 and Arg657, both of which are involved in the LRRLL motif. It is possible that phosphorylated Thr654 interacts with these and other Arg residues, resulting in disruption of the electrostatic interactions with acidic lipids.

In conclusion, based on our spFRET analysis combined with the nanodisc technique, we proposed a mechanism by which membrane lipid compositions dynamically regulate the formation and dissociation of TM-JM dimer in cooperation with protein phosphorylation. Acidic membrane lipids and Thr phosphorylation regulate the monomer/dimer equilibrium of JM-A by changing the interaction of the JM-A region and the membrane. In the cells, EGFRs are concentrated in the lipid raft with high concentration of PIP_2_^36^ and activate a variety of intracellular signal proteins^37^, including PKC and PLC-γ, upon ligand association. PKC phosphorylates Thr654 on JM-A domain^18^, and PLC-γ hydrolyzes PIP_2_ molecules in the cell membrane. Therefore, it is expected that although JM-PIP_2_ interactions help the dimerization of EGFR protomers when ligands bind to the extracellular domains, dimerized EGFR subsequently activates PKC and PLC-γ in the cytoplasm, resulting in the electrostatic perturbation of JM-PIP_2_ interactions and dissociation of EGFR dimers (Fig. 5d). Further analysis would provide a better understanding of the relationship between membrane lipid molecules and the activation of EGFR as well as other membrane receptors.

## Acknowledgments

The study was supported by the Japanese Ministry of Education, Culture, Sports, Science and Technology (MEXT) Japan (JP16H00788, JP15H02394, JP15KT0087).

## Author Contributions

R.M. performed sample preparation, spectroscopic assay, and single molecule fluorescence measurements. T.S. synthesized and purified fluorophore-labelled TM-JM peptides. R.M., K.O., M.Y., and Y.S. analyzed data. T.S. advised on experiments and manuscript preparation. R.M. and Y.S. wrote the manuscript with advice from all authors.

## Online Methods

### Materials

1-Palmitoyl-2-oleocyl-*sn*-phosphatidylcholine (POPC, PC), 1-palmitoyl-2-oleoyl-*sn*-phosphatidylserine (POPS, PS), and phosphatidylinositol 4,5-bisphosphate (PIP_2_) were purchased from Avanti Polar Lipids as chloroform solutions (PC and PS) or powders (PIP_2_). Cy3-maleimide and Cy5-maleimide were purchased from GE Healthcare Life Sciences. The detergent, *n*-octyl-β-D-glucoside (OG), was purchased from Dojindo. Monofunctional polyethylene glycol-succinimidyl valerate (PEG, MW 5000) and biotinylated monofunctional polyethylene glycol-succinimidyl valerate (b-PEG, MW 5000) were purchased from Laysan Bio, Inc.

### Peptide Synthesis and Purification

Peptides corresponding to the TM-JM regions of EGFR (618-666) were synthesized by solid-phase methods with the following sequence: KIPS-IATGMVGALLLLLVVALGIGLFM-RRRHIVRKRT_654_LRRLLQERELVE-C-NH_2_^29^. The synthetic peptides were purified by reverse-phase HPLC on a C4 column with a gradient of 1-propanol and acetonitrile (1:1) containing 0.1% aqueous trifluoroacetic acid (TFA). For fluorescent labeling of peptides, Cy3-maleimide or Cy5-maleimide was introduced to the sulfide group on cysteine at the C terminus of the TM-JM peptide by mixing the peptide and fluorescence derivative in dimethyl formamide under basic conditions.

### Nanodisc Preparation

For nanodisc preparation, the fluorescent EGFR TM-JM peptides cosolubilized with lipids and OG in hexafluoroisopropanol were dried to form thin firms. The peptide films were resolubilized in 0.5 M NaCl, 20 mM Tris, 0.5 mM EDTA, 30 mM OG, and 5 mM DTT (pH 7.5). His8-tagged membrane scaffold protein 1E3D1 (MSP1E3D1) was expressed in *E. coli* and purified as described previously^27^. The concentration of MSP was quantified based on the absorbance at 280 nm (29910 m^−1^ cm^−1^). Thin PC or PS films were formed by evaporation of the solvent (chloroform) under a stream of nitrogen gas and dried in vacuum. PIP_2_ powders were first dissolved in a solution containing chloroform, methanol, and water (20:9:1, v/v), and a thin film was formed as described above. PC, PS, and PIP_2_ were resuspended in a solution containing 0.5 M NaCl, 20 mM Tris/HCl, 0.5 mM EDTA, and 0.4 M sodium cholate (pH 7.5) at a final concentration of 10 mM. TMJM-Cy3 and TMJM-Cy5 were mixed in equal amounts and then conjugated with MSP and phospholipids (PC/PS) at a molar ratio of 1:1:120 μM (TM-JM:MSP:lipids). The mixture was dialyzed against Dialysis Buffer (Buffer D; 0.5 M NaCl, 20 mM Tris/HCl, 5 mM EDTA, pH 7.5) at 4°C to reconstitute the nanodiscs by removing detergent. The aggregates and liposomes were removed from the mixture by size exclusion chromatography using an ENrich™ SEC 650 column (Bio-Rad). PIP_2_ resuspended vesicles were added to the nanodisc solutions at the final concentration of 2 μM.

### Spectroscopic measurements

Fluorescence spectra were recorded with a Shimadzu RF-5300PC spectrofluorometer. Cy3 was excited at 540 nm and emission was detected through a glass cut-off filter (O57; Toshiba).

### Single pair FRET (spFRET) measurements

For single-molecule observation, nanodisc samples were immobilized on a glass surface as described previously^30,31^. Briefly, amine-modified glass surfaces were coated with 99% PEG and 1% b-PEG, and NeutrAvidin (Thermo Fisher Scientific) was then bound to b-PEG. The nanodisc samples bound with biotinylated anti-His8-tag antibody (MBL Life Science) were loaded into the glass chamber and allowed to bind to the NeutrAvidin-coated glass surface, after which unbound nanodiscs were washed away. To reduce the photobleaching rate of Cy3 or Cy5, the loaded nanodisc samples were filled with Buffer D containing 2-mercaptoethanol and 6-hidroxy-2,5,7,8-tetramethylchroman-2-carboxylic acid (Trolox) at final concentrations of 0.5% (w/v) and 1 mM, respectively. Fluorescence of Cy3 and Cy5 was observed using a TIRF microscope^22,38^ based on an inverted microscope (TE 2000; Nikon) with a 60× oil-immersion objective (ApoTIRF 60× 1.49 NA; Nikon). Fluorescence of Cy3 was generated using a 532 nm laser. Dual-color imaging was carried out through a 4.0× relay lens using EMCCD cameras (C9100-134, ImagEM; Hamamatsu Photonics) with 200× EM gain. Images of 512 × 512 pixels (67 nm/pixel) were recorded with a temporal resolution of 100 ms/frame.

### Analysis of FRET efficiency distributions

Analysis of TIRF images was performed using ImageJ as described previously^22^. The background noise was filtered using the *Subtract Background* function in ImageJ. Fluorescence intensities of Cy3 and Cy5 in single nanodiscs were measured as averages in circles with a diameter of 12 pixels containing a fluorescence spot. The average intensity of the same size of circles in which no spot was present was subtracted as the background. Along the fluorescence trajectories of TMJM-Cy3 and TMJM-Cy5, FRET efficiency, *E*_FRET_, for each frame was calculated from the fluorescence intensities in the donor *I*_D_ and acceptor *I*_A_ channels as

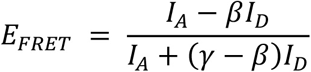

where *β* and *γ* are coefficients for compensation of fluorescence leakage from the donor dye to the acceptor detector channel and difference in detection efficiencies of dyes, respectively^23^.

### Bayesian change point detection

For dynamic analysis of FRET transitions, change points between different FRET efficiency states were determined using a modified Bayesian change point detection algorithm, as reported previously^34^. Briefly, we assumed two hypotheses; *H*_1_: A point *X* in the trajectory is not a change point in the vicinity of 2*N* frames. Points involved in *N* frames before and after point *X* follow the same distribution. *H*2: A point *X* in the trajectory is a change point in the vicinity of 2*N* frames. Points involved in *N* frames before and after point *X* follow different distributions.

Assuming that the FRET trajectory of each spot follows a Gaussian distribution, the likelihood for each of the hypotheses *P*(*D*|*H*_1_) or *P*(*D*|*H*_2_) is obtained and the ratio of two likelihood is defined as Bayes factor, as described previously^34^. When the Bayes factor overcomes a critical value (*B*), we consider point *X* as a change point. In addition, to eliminate falsely detected points, such as shot noise, a threshold of clean-up *C* is introduced, which integrates the traces if the lifetime of detected states is below *C*. The optimal parameters of *B*, *N*, and *C* were determined by comparing the numbers of detected change points (Fig. S4). We modified the original method^34^ to use a fixed value of *N* to prevent divergence in calculation.

